# Distinctions among electroconvulsion- and proconvulsant-induced seizure discharges and native motor patterns during flight and grooming: Quantitative spike pattern analysis in *Drosophila* flight muscles

**DOI:** 10.1101/481234

**Authors:** Jisue Lee, Atulya Iyengar, Chun-Fang Wu

**Affiliations:** Department of Biology, University of Iowa, Iowa City, IA, 52242, United States; Interdisciplinary Graduate Program in Neuroscience, University of Iowa, Iowa City, IA, 52242, United States

**Keywords:** Dorsal Longitudinal Muscle, Poincarè Plot, Instantaneous Firing Frequency, Instantaneous Coefficient of Variation

## Abstract

In *Drosophila*, high-frequency electrical stimulation across the brain triggers a highly stereotypic repertoire of spasms known as electroconvulsive seizures (ECS). The distinctive ECS spiking discharges manifest across the nervous system and can be stably assessed throughout the seizure repertoire in the large indirect flight muscles (DLMs). ECS discharges in DLMs have been extensively used to monitor seizure activities, notably in stress (‘bang’)-sensitive mutants. However, the relationships between ECS-spike patterns and native motor programs, including flight and grooming, are not known and their similarities and distinctions remain to be characterized. We employed quantitative spike pattern analyses including: 1) overall firing frequency, 2) spike timing between contralateral fibers, and 3) short-term variability in spike interval regularity (CV_2_) and instantaneous firing frequency (ISI^−1^) to reveal distinctions amongst ECS discharges, flight and grooming motor patterns. We then examined DLM firing phenotypes in well-established mutants in excitatory cholinergic (*Cha*), inhibitory GABAergic (*Rdl*) and electrical (*ShakB*) synaptic transmission. The results provide an initial glimpse on the vulnerability of individual motor patterns to perturbations of respective synaptic transmission systems. We found marked alterations of ECS discharge spike patterns in terms of either seizure threshold, spike frequency or spiking regularity. In contrast, no gross alterations during grooming and only a minor reduction of firing frequency during Rdl mutant flight were observed, suggesting a role for GABAergic modulation of flight motor programs. Picrotoxin (PTX), a known pro-convulsant that inhibits GABA_A_ receptors, induced DLM seizure patterns that displayed some features, e.g. left-right coordination and ISI^−1^ range, that could be found in flight or grooming, but distinct from ECS discharges. Our results indicate that these quantitative techniques may be employed to reveal overlooked relationships among aberrant motor patterns and native DLM motor programs in genetic and pharmacological analyses of underlying cellular and neural circuit function.

## Introduction

For nearly half a century, *Drosophila* mutants have opened the avenue to the mechanistic elucidation of how genetic variations contribute to the physiological basis of epileptiform behaviors and related seizure disorders [1, 2, 3]. Early mutants were isolated on the basis of seizures triggered by stressors such as high temperature (e.g. *shibire* [4], *seizure* [5]) or mechanical shock (e.g. *bang-senseless, easily shocked, knockdown*, and *bang-sensitive* [6]), while other mutants displayed spontaneous spasms (e.g. *Shudderer* [7]). Mutational studies have led to identifications of individual genes associated with seizure disorders, including those encoding for or regulating ion channels [8, 9, 10, 11], ionic transport and distribution [12, 13, 14], and enzymes related to energy metabolism [15, 16, 17, 18]. Different adult preparations [19, 20, 21, 22] have been employed to monitor seizure activity in the nervous system. In particular, the indirect flight muscles, dorsal longitudinal muscles (DLMs), have been most widely utilized as a readout of aberrant motor activities, especially during seizure discharges. High-frequency stimulation across the brain can trigger an electroconvulsive seizure (ECS [20]) with a highly stereotypic repertoire consisting of an initial spike discharge (ID), a period of paralysis, and a second delayed spike discharge (DD [21]). This seizure repertoire is modified in specific manners characteristic of individual seizure mutants [21, 22]. This line of studies demonstrates that the large DLM can provide convenient but reliable readouts for epileptiform and other patterned neural activities generated in the central nervous system (see [20, 23, 21, 22, 24, 25]).

A number of activity patterns driven by the innervating motor neuron (DLMn) have been characterized in detail, including flight, grooming, courtship and giant-fiber mediated jump-and-flight escape. During flight, the DLM fibers (six on each side) undergo myogenic, stretch-activated, isometric contractions to generate tension in phase with the oscillation of thorax case to power wing depression during the wing beat cycles (~ 200 Hz, [26, 27]). DLM action potentials, evoked by DLM motor neuron (DLMn) spikes, fire continuously and rhythmically at approximately 5 Hz [28, 29] solely to facilitate Ca^2^+ influx required for continued oscillation in muscle tension [30, 31]. In contrast, discrete individual spikes are associated with giant-fiber pathway activation [32, 33, 34, 35, 36], while distinct spike bursting activities are associated with grooming [34] and courtship song production [37].

In this report, we quantitatively describe the firing patterns of the DLM during electroconvulsion-induced seizure discharges versus the native spiking patterns during flight and grooming. In addition to ensemble statistics, we analyzed bilateral coordination of motor activities, as well as the local temporal features of spike trains, including instantaneous firing rate, short-term firing regularity, and timing relationships among adjacent spikes.

We further examined how activity patterns characteristic of each motor programs were affected upon genetic or pharmacological manipulations. We studied well-characterized mutants of major neurotransmission systems known to have clear alterations in physiology and behavioral expression: 1) picrotoxin (PTX), a non-competitive antagonist of the GABA_A_ receptor [38]; 2) a related receptor mutant, *Resistant to dield-rin (Rdl)* that encodes the GABA_A_ receptor α subunit [39]; 3) *Choline acetyltransferase (Cha)* mutants with reduced acetylcholine (ACh) synthesis [40, 41]; and 4) mutants of *ShakB*, encoding an innexin gap junction protein required for transmission at electrical synapses [42]. These treatments revealed clear quantitative distinctions between the seizure discharge patterns evoked by ECS and that induced by GABA receptor blockade via PTX. Thus, the quantitative approaches undertaken here may be applicable to the study of additional aberrant DLM spike discharges to explore their potential association with or distinction from the endogenous spike patterns during normal motor activities.

## Methods

### Fly Strains

*Drosophila melanogaster* strains were kept in standard glass vials containing cornmeal medium [43] and reared at room temperature (23 °C). Adult flies used for the experiments were 3-12 days old. Wild-type flies were of the strain *Canton S* (CS). Synaptic transmission mutants examined include those alleles defective in *Cha* encoding a Choline Acetyltransferase *(Cha^ts2^* [44]), *Rdl* encoding the GABA_A_ receptor a subunit (*Rdl^MD-RR^*, [39]), and *ShakB* encoding the structural subunit of an innexin *(ShakB^2^* [45]; *ShakB^Pas^* [42], also known as *ShakB^3^*, [46]). Data from male and female individuals were pooled, as previous results have indicated that DLM firing properties are not generally different between sexes during flight or seizure activity [21, 22, 27].

### Tethered Preparation and DLM Recording

The fly tethering procedure used for DLM recordings have been described in detail previously [21, 22, 27]. Flies were briefly cold- or ether-anesthetized and glued to a tungsten pin using cyanoacrylate or nitrocellulose based glue. Flies were allowed to recover from anesthesia for at least 30 min prior to recording. All recordings were done at room temperature (~ 23 ^°^C).

Electrolytically sharpened tungsten electrodes were used to access DLM spikes, with an electrode inserted into the dorsal-most fiber (DLMa, 45a, c.f. [47]), and a similarly constructed electrode inserted into the abdomen for reference. Signals were amplified by an AC amplifier (either DAM-5A, World Precision Instruments, New Haven CT; or AM Systems Model 1800, Carlsborg, WA) and digitized via a data acquisition card at a sampling rate >10 kHz (Digidata 1200, Axon Instruments, or USB 6210, National Instruments). Spike detection and analysis were done in PClamp 6, LabVIEW 8.6 and in MATLAB r2017b using custom-written scripts.

### Electroconvulsive Stimulation

Electroconvulsive stimulation was delivered across the brain via tungsten electrodes inserted into each eye, and consisted of a train of 0.1-ms pulses delivered at 200 Hz at a specified voltage (30 – 100 V) and train duration (0.5 – 4 s). Seizure susceptibility of individual flies was examined for each genotype (Figure 6) using five different intensity levels 1 – 5 (stimulus voltage, train duration): 1. (50 V, 0.5 s); 2. (50 V, 1.0 s); 3. (50 V, 2.0 s); 4. (100 V, 2.0 s); and 5. (100 V 4.0 s). Each level corresponds to a progressive doubling of stimulation intensity [22]. When appropriate, after the electroconvulsive stimulation, test pulses sufficient to directly activate the giant-fiber neuron (24 V or higher, 0.1 ms duration, delivered at 1 Hz) were applied to examine synaptic transmission along giant-fiber pathway.

### Pharmacology

Picrotoxin (PTX) was used to block ionotropic GABAergic signaling [21]. Two methods were employed to administer PTX systemically in intact, tethered flies: 1) small droplet applied on an eye incision (‘eye drop’), and 2) a fixed volume injection into the dorsal vessel (‘DV inejection’). For eye drop application, a stimulation electrode was advanced to penetrate the basement membrane of the retina and then retracted. A small drop of 1mM PTX solution dissolved in 0.6 % NaCl solution (or Control NaCl solution without PTX) was applied to the incision and was allowed to diffuse to the hemolymph. The DV injection procedure was adapted from Howlett & Tanouye [48]. A filamented glass electrode (1.0 mm OD, 0.58 mm ID, A-M Systems #601500) was pulled in a Brown-Flaming electrode puller (P87, Sutter Instruments, Novato, CA). After loading the distal end of the electrode with PTX solution, capillary action consistently drew 0.33 *μ*l of solution to the tip. To inject the loaded solution, the electrode tip was broken and was inserted into the dorsal vessel [47] of the tethered fly. The solution was then injected using air pressure delivered manually through a syringe.

### Spike Train Analysis

Firing rate was analyzed using two measures: instantaneous firing frequency and overall spiking rate over a specified time window of interest. The instantaneous firing frequency was defined as the reciprocal of the inter-spike interval (ISI) measured in seconds, between successive spikes [49, 50], referred to as ISI^−1^ with units of Hz. For sequential ISI^−1^s in a spike train, the occurrence of the first spike is used to mark the temporal location of the particular inter-spike interval. The overall spike rate was defined as the total spike count during the specified time window divided by its duration. Poincaré trajectories (Figure 4, Figure 7 and Figure 9) were constructed by plotting the time series of ISI^−1^ within a spike train, with each ISI^−1^ against the ISI^−1^ of the subsequent interval, i.e. ISI^−1^_*i*_ vs. ISI^−1^_*i*+1_. Therefore for a spike train of *n* spikes, there will be *n* – 1 data points in the Poincaré trajectory.

To quantify the regularity of ISIs within a spike train, the coefficient of variation, CV, defined as the standard deviation of ISIs divided by the mean ISI, has been extensively used (e.g. [51]). However, to accurately estimate variability, the average firing frequency of the spike train is usually assumed to be stationary over a sufficiently long period of firing for which the CV is computed. To assess variability in spike trains with varying firing frequency over relatively short time periods, Holt and associates [52] developed CV_2_ which estimates variability between adjacent spike intervals. The CV_2_ is computed for each spike sequentially along the spike train from each pair of two adjacent intervals ISI_*i*_ and ISI_*i*+1_ as:

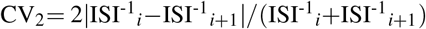

For cases of spike trains with stationary average firing rates over time, estimates of variability derived from CV and CV_2_ are in close agreement [52]. For each spike train, the scatterplot of the ISI^−1^ versus CV_2_ documents the local firing frequency (or ISI-1), and firing frequency variability (CV2) around individual spikes (represented as points in the 2-D plot). After connecting temporally adjacent spike-points, the trajectory thus traces the history of the spike-interval variability along with the sequential instantaneous spike firing frequencies. Note that the plot of the ISI^−1^ versus CV_2_, may be derived from the Poincaré trajectory, which registers sequentially the ISI^−1^ before and after each individual spike throughout the same spike train.

For assessment of the general trends and for clarity in presentation, we applied a running average filter to smoothen the raw ISI^−1^ vs. CV_2_ plots. We verified that filter window sizes ranging from 3-9 pts provided qualitatively similar results without severely distorting major features of the trajectory for the three motor programs examined here. A 6-spike window was applied in ISI^−1^ vs. CV_2_ plots because it appropriately captures the characteristics for both sustained flight and short grooming bouts (Supplemental Figure 1).

The filtered ISI^−1^ vs. CV_2_ plots also enabled us to obtain smoothened ‘overall’ trajectories for the different motor patterns based on the ensemble of spike trains from recorded individuals, even though each train could differ somewhat in numbers of spikes. Briefly, durations of individual spike trains were normalized, such that ‘0’ represented the initial spike and ‘1’ the last spike, with the relative temporal locations for the rest of spikes reassigned proportionally along the normalized unitary duration. Thus, an overall average trajectory of a specific motor pattern could be constructed based on the collective temporal information from an ensemble of spike trains, i.e. to determine the averaged ISI^−1^ vs. CV_2_ values, or (ISI^−1^, CV_2_), along the unitary duration. For a spike train in the ensemble, each rescaled ISI segment still carries the same (ISI^−1^, CV_2_), associated with the ith spike. The averaging process for the ensemble (ISI^−1^, CV_2_) was carried out with an increment of 0.01 (bin size) along the normalized duration. The resulting variation of (ISI^−1^, CV_2_) along the normalized the time base, or unitary duration, was re-plotted in the 2-D graph with ISI^−1^ vs. CV_2_ coordinates. The range of number of ISIs for this treatment was between ~60 and ~300 for DD, ~400 and ~1200 for flight, and 20 and ~500 for grooming spike trains. The outlier spike trains were eliminated from the calculation (mostly occurred in grooming).

The bilateral phase (*θ*) relation between individual spikes from the left and right DLM units was calculated as the proportion of the time point within the contralateral inter-spike interval that the ipsilateral unit generated a spike, i.e. *θ* ranges from 0 to 1 [28, 53].

## Results

### Three Identified Motor Programs Driving DLM Activity: Electroconvulsive Seizure Discharges, Flight and Grooming

High frequency stimulation across the brain triggers a previously described ECS repertoire in wild-type (WT) and different mutants [20, 21], which consists of an initial discharge (ID) of DLM spikes corresponding to an initial spasm, a period of paralysis where conduction fails along the giant fiber (GF) jump-and-flight escape pathway, followed by a delayed discharge (DD) of DLM spikes, and finally recovery of GF pathway transmission and voluntary motor behaviors. The present study focuses on DD spike patterns since quantitative aspects of DLM spiking during the ID have been characterized [17, 27] whereas spike patterning during the DD has received less attention [22]. We noted that flies often display a ‘wings-up’ posture during DD (Figure 1A, Supplemental Video 1, cf. [21]). During flight, the wings beat at approximately 200 Hz [54, 55, 27], and during grooming, wing depression facilitates leg sweeping across the wing (Figure 1A, [34, 56, 57]). We observed characteristic patterns of DLM spiking corresponding with these behaviors (Figure 1B), each with distinct profiles of instantaneous firing frequency (defined as the reciprocal of each inter-spike interval, ISI^−1^ with units Hz, Figure 1C). Electroconvulsion-induced DDs consisted of DLM activity of varying firing frequencies, which quickly accelerates, peaks (~ 20 Hz), and gradually decelerates (Figure 1C). During sustained flight, the DLM firing was more rhythmic, with highly regular spike intervals (~ 5-10 Hz), whereas DLM firing during grooming differed markedly, with short bursts of spikes (ISI^−1^ > 100 Hz) coincident with wing depression events.

**Figure 1.**
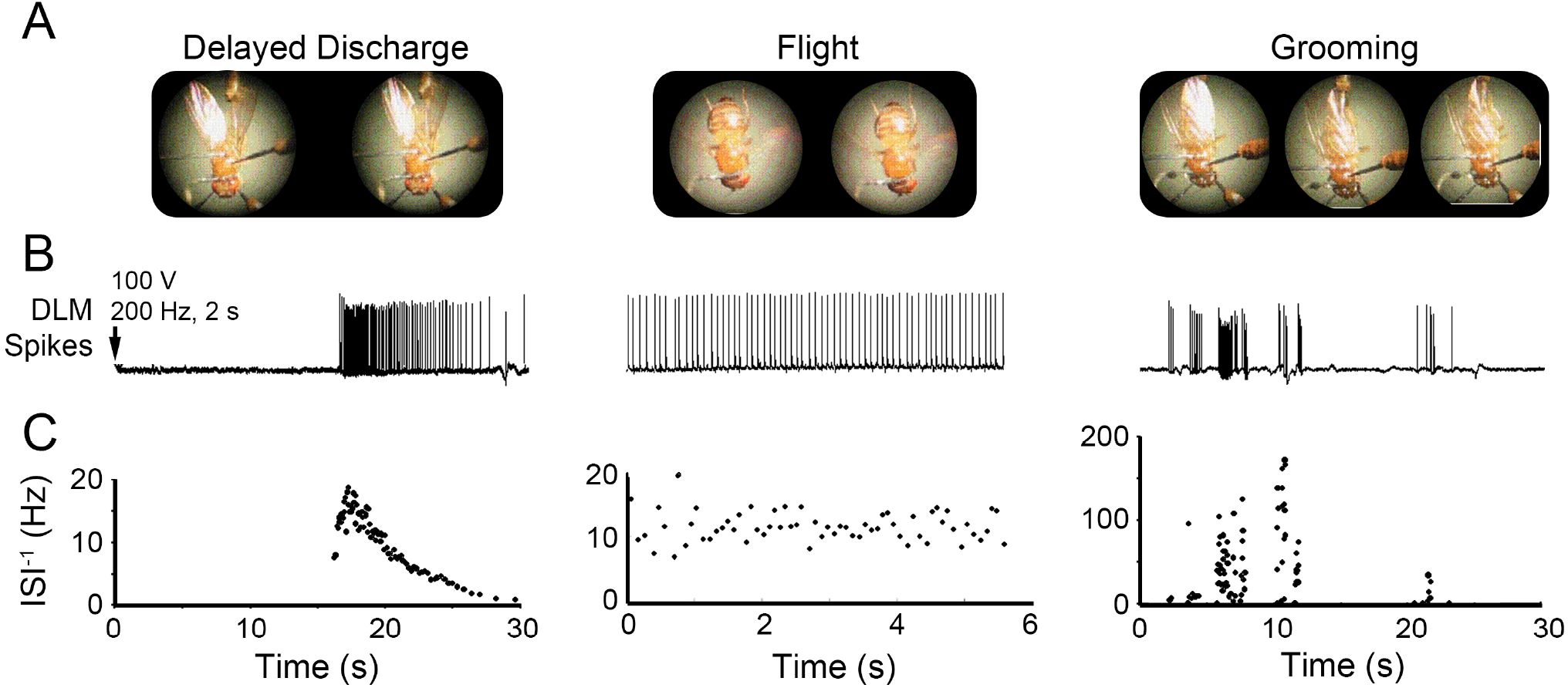
Wing posture and corresponding DLM spiking activity during delayed seizure discharges, flight and grooming. (A) Representative consecutive video frames (50 ms apart) of tethered fly behavior taken during the respective motor programs. During the delayed seizure discharges (DD), wings were often held in an ‘up’ position, although brief wing buzzes were sometimes observed. During flight the wings beat at approximately 200 Hz [27]. Grooming bouts were often characterized by the fly depressing its wing to enable leg sweeps over the dorsal wing surface. (B) Example traces of DLM spiking corresponding to the behaviors in (A). Arrow indicates timing of electroconvulsive stimulation triggering delayed discharges. Note the rhythmic DLM spiking activity during flight and bursts of activity associated with grooming behavior. (C) Plots of the instantaneous firing frequency (inverse of inter-spike intervals, ISI^−1^) versus time corresponding with the respective traces in (B). The ISI^−1^s observed during grooming bouts were an order of magnitude higher than firing observed during either DD or flight.

Figure 2A depicts histograms of the ISI^−1^s during the respective motor programs from a representative fly. It is evident that the three motor programs had clearly distinct frequency profiles. DD seizure spiking displayed a right-skewed ISI^−1^ distribution with most spiking occurring between 1 and 10 Hz, with a clear “tail” of vanishing activity and a briefer session of fast-firing spikes, beyond 25 Hz, corresponding with the early peak of DD firing. During flight, the distribution of ISI^−1^s was markedly less variable, reflecting a rhythmic firing. During grooming, DLM spiking displayed a highly-variable ISI^−1^ distribution, with some short spike bursts exceeding 100 Hz—well above the maximum rate observed during either DD or flight.

**Figure 2.**
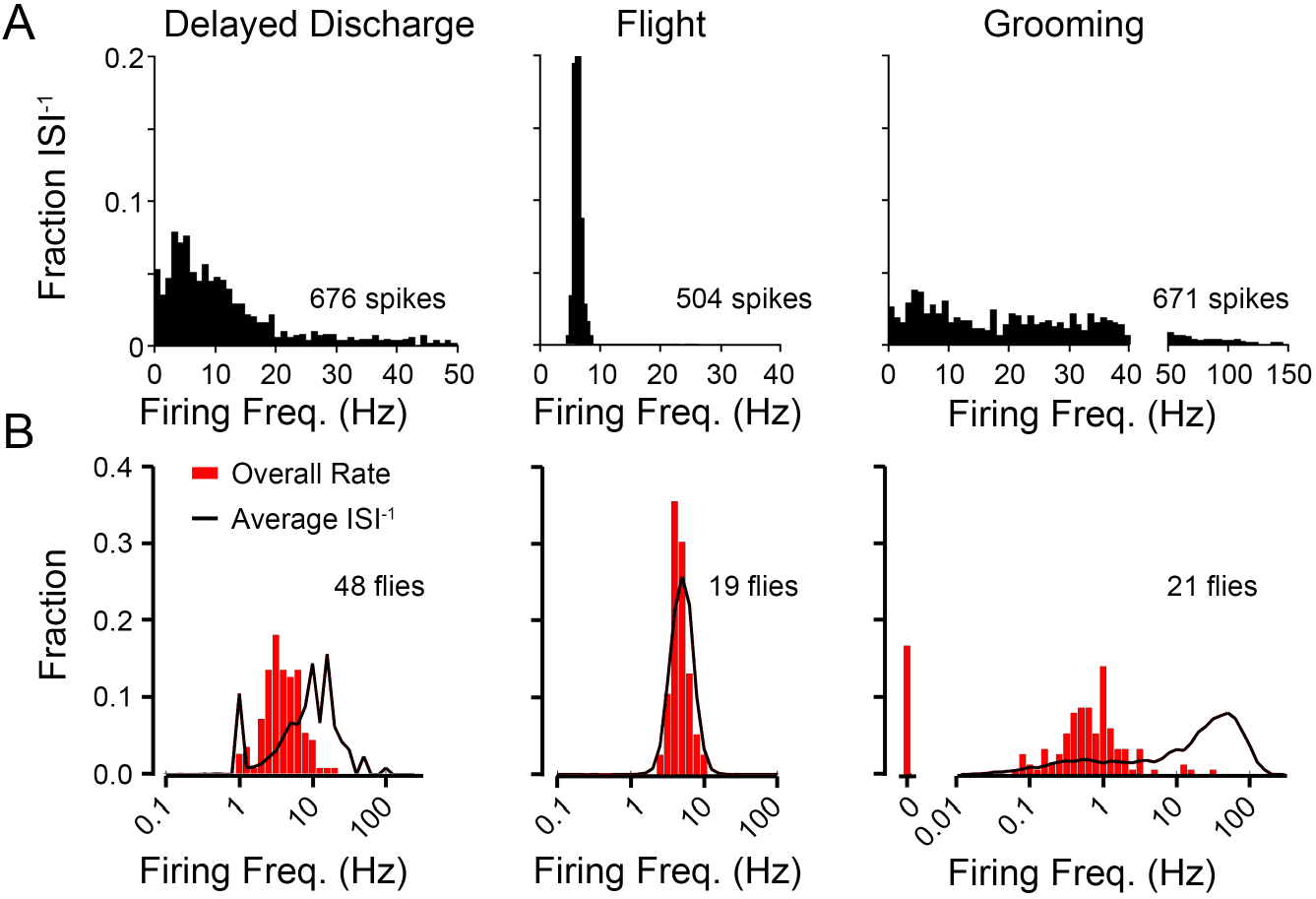
DLM firing characteristics during delayed seizure discharge, flight and grooming activities. (A) Histograms of DLM instantaneous firing frequencies observed during DD, flight and grooming bouts from representative individuals. The DD histogram was constructed from nine discharges evoked in a single fly (total of 92 s of activity, 676 spikes), flight and grooming histograms were obtained from 130 s bouts of the respective behaviors (1583 and 671 spikes respectively). (B) Distributions of two measures of firing frequency during respective motor programs across a population of WT flies. Red histogram, the average of the overall firing frequency, defined as the total spike count/total recording duration for each spike train. Black line profile, the ensemble distribution of instantaneous firing frequencies (ISI^−1^) from the same spike trains. Note that the distributions are plotted on a log-frequency scale. Sample sizes as indicated.

With an ensemble of tens of flies, we examined firing rate regularity across the population for each motor pattern, by comparing the fractional distribution of the average ISI^−1^ along the spike train against that of the overall firing rate (defined as the total spike count divided by recording duration, typically 60 – 120 sec). In general, rhythmic firing produces similar results from the two measures and disparity between the two measures suggests highly arrhythmic firing. As Figure 2B illustrates, rhythmic DLM firing during flight produced highly similar distributions of firing rate based on the two measures (average ISI^−1^ = 6.70 Hz, overall firing rate = 4.90 Hz), whereas grooming behavior was associated with irregular short bursts of a few spikes during, leading to drastically different rate distributions derived from the total spike count or from the instantaneous firing frequency (average ISI^−1^ vs. overall firing rate = 16.62 vs. 0.55 Hz). The discrepancy between the two measures of firing rate during delayed seizure discharges fell in between (average ISI^−1^ vs. overall firing rate = 7.46 vs. 5.08 Hz).

### Interactions between Bilateral DLM Motor Units during Delayed Seizure Discharges, Flight and Grooming

Analysis of spike timing relationships between left and right DLM pairs can reveal potential interactions between the bilaterally symmetrical pairs of motor neurons that innervate them and can yield important clues to distinctive mechanisms of spike pattern generation among different motor programs. We recorded from top-most pair of DLMs (#45a [47]), each of which is innervated by a contralateral MN5 motor neuron in the mesothoracic ganglion [58]. Of the three motor patterns examined here, relationships between bilateral DLM pairs during flight have been best characterized [28, 29, 53], while the relationships during grooming and delayed seizure discharges have yet to be documented. Within individual flies, we found that for flight and DD, the firing frequency profiles of the left and right DLM units were not identical, but similar to each other (Figure 3A, top two panels, data from a representative individual). In contrast, striking discrepancies in firing frequency between left-right DLM pairs were observed during grooming (Figure 3A, bottom panel). In behavioral context, the left-right pairs perform coordinately during flight and DD, but can be decoupled during grooming.

**Figure 3.**
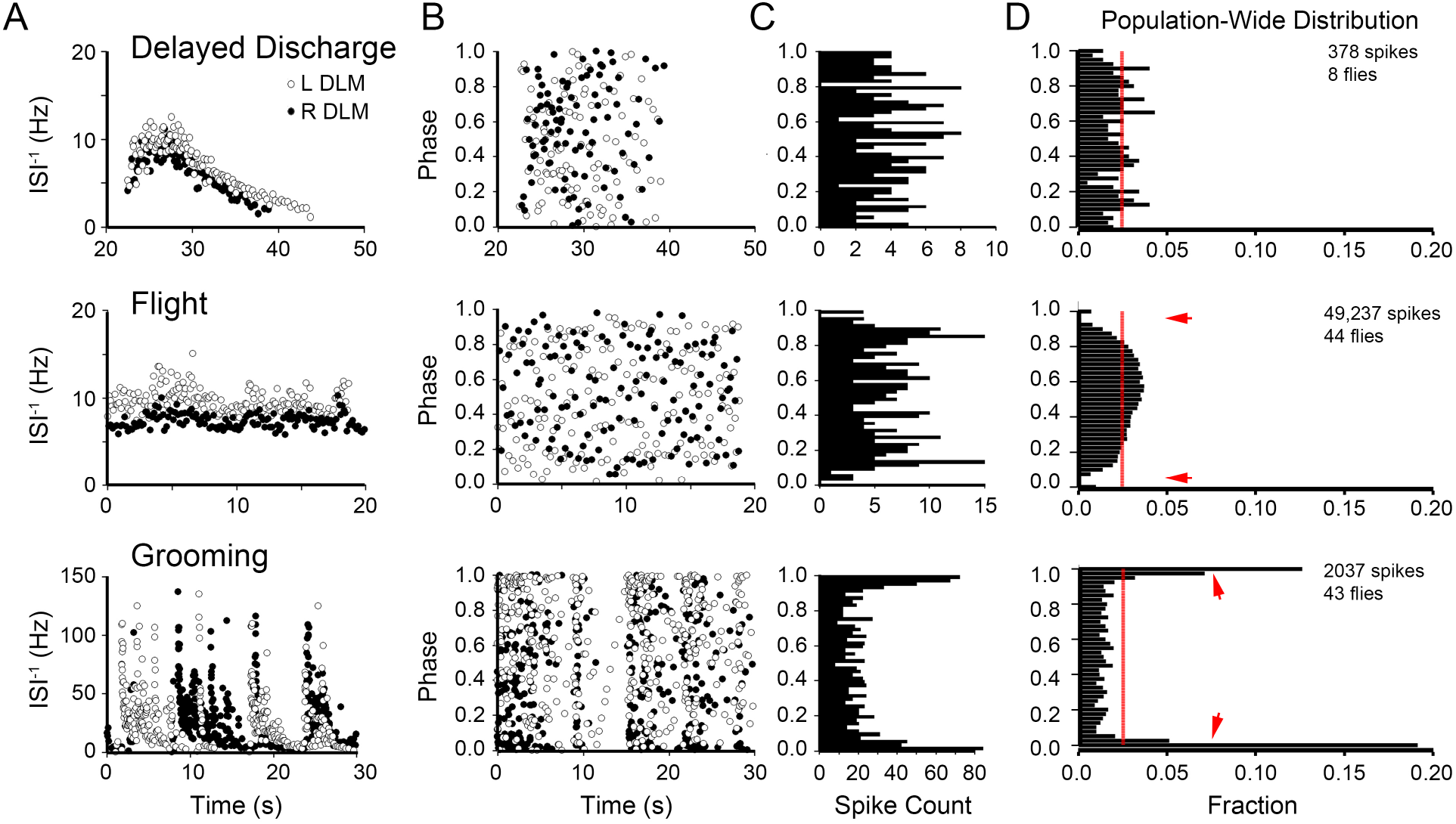
Timing relationships between left and right DLM spikes during delayed seizure discharges, flight and grooming. (A) Plots of the instantaneous firing frequency time course for left (open circles) and right (closed circles) DLM spiking during DD (top panel), flight (middle panel) and grooming behavior (lower panel) from a representative individual. (B) Corresponding plots of the phase relation between left and right DLM spiking for data shown in panel A. The bilateral phase relation between left and right DLM spikes was determined as the fraction within the inter-spike interval from the contralateral unit that lapsed prior to the ipsilateral spike occurrence [28, 53]. Open circles indicate the phase location of the left DLM spikes with respect to the right DLM spiking, and vice-versa for closed circles. (C) Histograms of the bilateral phase relations from the right DLM spikes of the respective plots in B (closed circles). (D) Ensemble histograms of the bilateral phase relationships from the population of flies observed during the three motor programs, across the population of flies observed (sample sizes indicated). Red line indicates a uniform phase distribution. Arrows indicate phase-exclusion observed during flight and in-phase firing during grooming activity.

We further quantified bilateral phase relationship of DLM firing in the three motor programs by using the protocol described by Koenig & Ikeda [53]. Briefly, the fraction of an inter-spike interval that elapsed prior to the contralateral muscle firing was measured to define the left to right (or right-to-left) phase from zero to one for individual spikes along the left (or right) DLM spike train (Figure 3B, data from a representative individual, see also Methods). Histograms of the bilateral phase relationships between contralateral units appeared relatively unremarkable during DD, compared to the other two motor programs (Figure 3C, derived from Figure 3B). An ensemble distribution created from DD spike trains across different flies did not indicate significant deviation from a uniform distribution (Figure 3D, p > 0.05, *χ*^2^-test). However, during flight, the bilateral phase relations of DLM units displayed a characteristic spike exclusion, as indicated by the observation that immediately after a spike, the contralateral unit rarely spiked (± 10% of ISI period, approximately 40 ms; arrows in Figure 3D). This “exclusion band” feature was previously described by Levine & Wyman [28], and may serve to differentiate flight-associated spiking from DD seizures within individual DLMs. In contrast to the relatively uniform distributions associated with DD and bilateral exclusion during flight, grooming spike activity was characterized by phase-correlated firing in the left-right DLM pairs, with most firing occurring within 10% of the ISI period on the contralateral side. Thus, our analysis of the phase relationship provides a quantitative indicator for degrees of independence, exclusion or coupling between bilateral pairs of DLMs during the respective motor programs.

### Poincaré and ISI^−1^ vs CV_2_ Plots: Distinct Signatures of Rate and Rhythmicity among DLM Spike Patterns

In addition to the ensemble ISI^−1^ distributions in a somewhat overlapping ranges (Figure 2), DD, flight and grooming spike patterns may be qualitatively distinguished based on differences in variation of ISI^−1^ between adjacent spikes (Figures 1 & 3A). The temporal structure of sequential spike intervals may be distinct among these firing patterns, but are not retained in the firing frequency distributions and left-right phase relationships (Figures 2 & 3B, C). Poincare plots (also known as return maps, c.f. [59, 60]) have been used to visualize spike interval dynamics, which may contain both deterministic and stochastic components. We plotted sequentially the ISI-1each of spike interval against the subsequent interval (ISI^−1^_*n*_ vs. ISI^−1^_*n*+1_, hereafter designated as Poincaré trajectory or PT), to quantify sequential changes along the progression of a spike train (Figure 4A). When plotted in this manner, a constant-frequency spiking train will have successive points in the PT falling at the same location on the lower-left to upper right diagonal. A spike train with gradual frequency-modulation will have data points wandering along the diagonal of the plot reflecting changes in the ISI^−1^. Irregular spike trains will have data points deviating from the diagonal to different degrees depending on how abruptly the sequential ISI^−1^ changes. Within individual flies, grooming spike patterns displayed much higher variability in successive spike intervals in their PT, compared to both DD and flight patterns that showed lower ISI^−1^ to ISI^−1^ variability. Thus, the PT for individual grooming spike trains traversed abruptly in broad strides across the diagonal, whereas successive data points in DD clustered closer to the diagonal and flight to almost a point (Figure 4A), uncovering the special morphological features of the spike trains hidden in the above population-level firing frequency distributions (Figure 2B).

**Figure 4.**
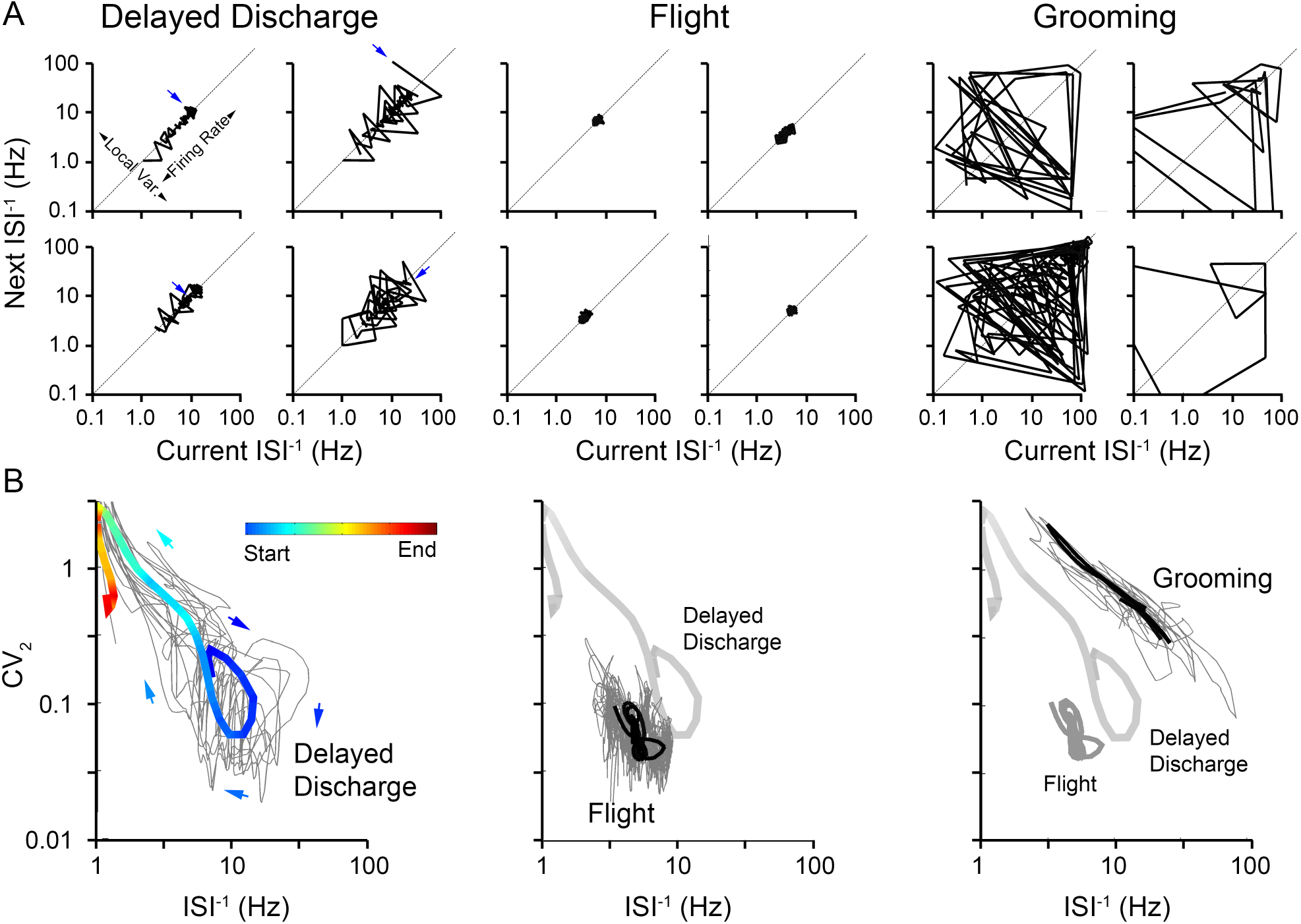
Poincaré plots and ISI^−1^ vs. CV_2_ trajectories of DLM spiking during delayed seizure discharges, flight and grooming. (A) Representative Poincare plots of DD, flight and grooming spike trains. For each inter-spike interval within the spike train, the ISI^−1^ is plotted against the subsequent ISI^−1^ within the sequence. Points along the lower-left to upper-right diagonal represent stable firing frequencies while deviations from the diagonal represent degrees of change in firing frequency. Arrows indicate start of trajectories for DDs. Note that axes are plotted on a logarithmic ISI^−1^ scale. Note the compact Poincaré plots of flight and the large deviations from the diagonal within the grooming plots. (B) Trajectories of the ISI^−1^ versus instantaneous coefficient of variation (CV_2_) for spiking during the three motor programs. Lower values of CV_2_ indicate more rhythmic firing. Ten sample trajectories of DD (left), flight (center) or grooming (left) trajectories are displayed (light grey), on which an ensemble average of the individual trajectories for the respective motor programs is over-laid (bold line) (ensemble average traces drawn from a sample size of 11, 126, and 115 individuals respectively). For the averaged DD trajectory, the line color and arrows indicate the trajectory’s time course and the ensemble average trajectories for flight and grooming are bolded. For the flight panel, the average DD trajectory is displayed for reference (bold grey line). For grooming trajectories, the averaged DD and flight trajectories are displayed for comparison. See Methods for computation details.

The above Poincaré trajectories clearly illustrated distinctions among the three motor patterns. However, multiple PTs cannot be readily treated to produce an ensemble trajectory based on larger sample sizes to highlight the global dynamic differences between motor patterns. We employed a transformation of the PTs to enable over-lays of individual trajectories for construction of an ‘average’ trajectory across a number of individual trajectories (Figure 4B). It is known that the distributions of both instantaneous firing frequency and variability between successive spike intervals can be extracted directly from the Poincaré plot [52, 61]. Since deviations of the PT from the diagonal are related to variability of spike train intervals, one established treatment is to quantify the deviation from the diagonal along the local firing frequency adjacent to each spike in the sequence by adopting a measure for the instantaneous coefficient of variation (CV2), defined as 2 lIS^−1^_*n*_ - ISI^−1^_*n*+1_| / (ISI^−1^_*n*_ + ISI^−1^_*n*+1_) ([52] see also Methods). The transformed plot of the ISI^−1^ vs CV_2_, using the same sequential ISI^−1^ information, is essentially a re-scaling of the sequential deviations from the diagonal in the PT plotted against the ISI^−1^. After this transformation, high CV_2_ values correspond with abrupt, large changes in adjacent spike intervals in either direction, increasing or decreasing.

This treatment generates ISI^−1^ vs CV_2_ plots that display clear distinctions among the three motor programs with an advantage of generating ensemble statistics for repeated trials (see Methods for averaging details). During delayed seizure discharges (DD), average trajectory of ISI^−1^ vs CV_2_ (Figure 4B, left panel, temporally color coded, average trajectory superimposed on individual trials in grey lines) illustrates a stereotypic trend of an initial acceleration (ISI^−1^ from ~7 to ~15 Hz, blue segments) accompanied by increasing regularity, lower CV_2_ (< 0.1), which then gradually decreases in firing frequency as well as regularity with increasing CV2 values (Figure 4B, left panel, cyan-green segment, climbing towards the upper-left corner, cf. Figure 11C, left panel). For sustained flight, DLM spiking appears as a tight trajectory with relatively little drift within a confined region (Figure 4, middle panel, ISI^−1^ ~5 Hz, CV_2_ ~0.03, cf. Figure 1C middle panel). Notably, the initial phase of the DD trajectory and the flight trajectories occupy adjacent regions of low CV_2_ values; although for DD (thick grey average trajectory re-plotted in Figure 4B for comparison), instantaneous firing frequencies are somewhat higher (~ 10 – 15 Hz vs 3 – 10 Hz) across the population sampled. In contrast, for the case of grooming, DLM spike trajectories occupy a region separate from either DD or flight, displaying significantly higher CV_2_ values, usually above 0.5, consistent with the qualitative observations of arrhythmic firing punctuated with abrupt changes in ISI^−1^ (Figure 4B, right panel cf. Figure 1C right panel). Thus, the temporal evolution of the spike patterns of the three motor programs are intrinsically distinct since the above quantitative treatment generates averaged trajectories that occupy three separable regions in the ISI^−1^ vs. CV_2_ plot.

### Differential Vulnerability of Three Motor Patterns to Genetic Perturbations of Synaptic Transmission Systems

To provide a first glimpse of the general vulnerability of spike patterns of the three motor programs to perturbations of the major synaptic transmission systems, we examined mutants of three genes that have been well-established to regulate inhibitory, excitatory and electrical synapses in the CNS: *resistant to dieldrin (Rdl*, encoding the inhibitory GABA_A_ receptor subunit [39]), *choline acetyltransferase* (*Cha*, acetylcholine synthesis enzyme [40]) and *ShakingB* (*ShakB*, encoding innexin gap junction subunits[62, 63, 64]). These mutants have been well-characterized for their effects on the giant-fiber pathway, from the brain afferents to DLM output [41, 42, 21]. We specifically examined the *Rdl^MD-RR^* allele, a natural variant of Rdl that has developed resistance to the insectide dieldrin, which also displays enhanced GABAergic inhibition [65, 66, 67]. The *Cha* allele examined, *Cha^ts2^* displays defects in cholinergic transmission at room temperature, even though its enzymatic activity is more severely disrupted at elevated temperatures [40, 41]. Both *ShakB* alleles studied here (*ShakB^2^* and *ShakB^Pas^*) are known to severely compromise signal transmission along the GF pathway that involves identified electrical synapses [42, 68].

We first examined instantaneous firing frequency of the three motor patterns (cf. Figure 1). Notably, DD discharges in the respective mutants showed distinct defects (Figure 55A). A substantial reduction was observed in both the number of spikes and the peak instantaneous firing frequency in *Rdl* mutants with enhanced GABAA receptor function (mean peak ISI^−1^ of 6.4 vs 13.0 Hz for WT, Figure 5Aii). Interestingly, disruption of excitatory cholinergic transmission in *Cha* mutants also led to a reduction in instantaneous firing frequency (ISI^−1^, 6.3 Hz), comparable to *Rdl* mutants. In contrast, altered electrical transmission in *ShakB* mutants did not significantly alter the peak ISI^−1^ in the DD (Figure 5Aii). However, the qualitative temporal profile was more resistant to these genetic manipulations, since across the three mutants, the rate of rise to the peak firing frequency remained unaltered, despite the clear modifications in their firing frequencies, (Time to Peak, Figure 5Aiii).

**Figure 5.**
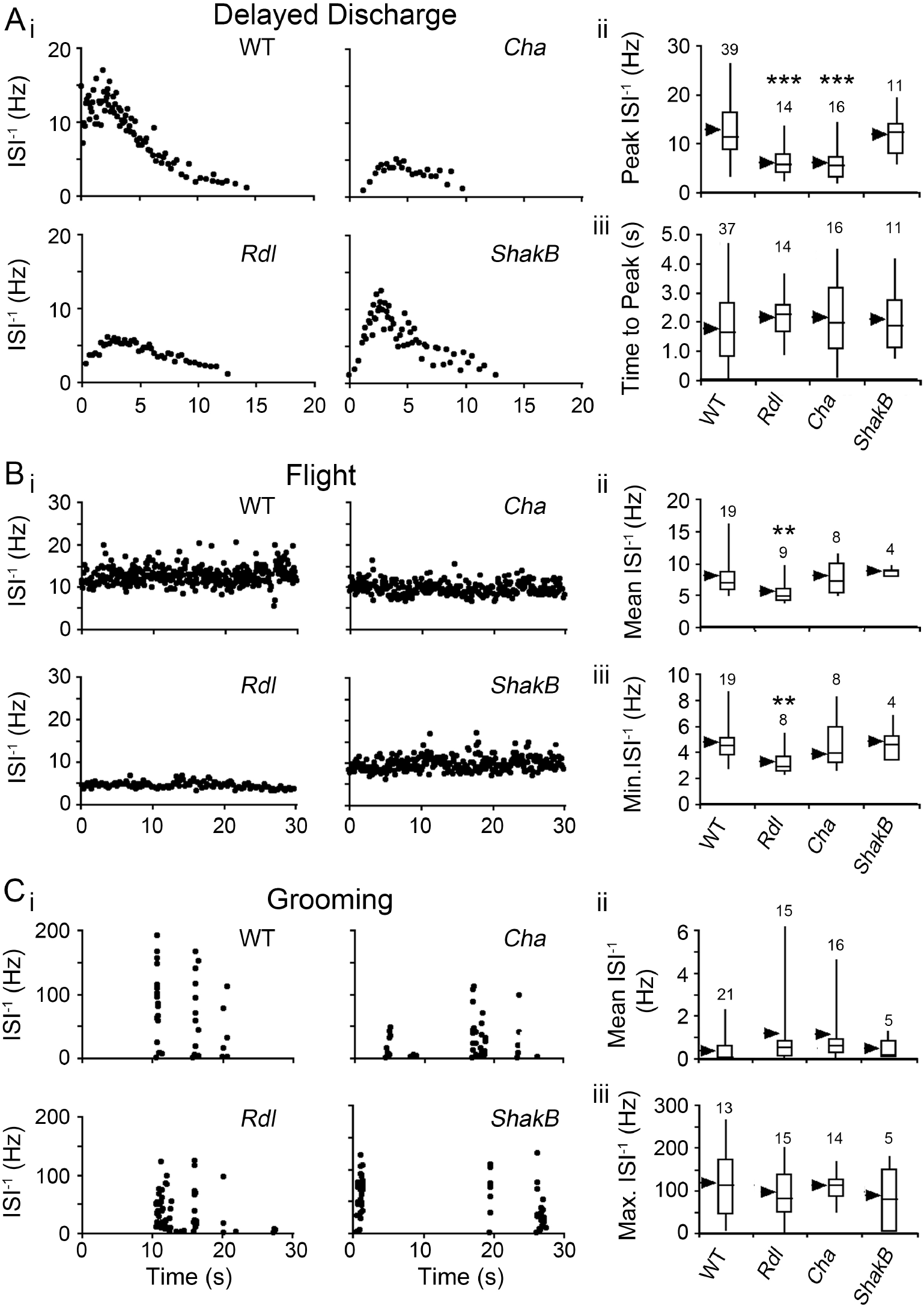
Modifications of DLM firing characteristics during DD, flight and grooming activity in WT, and the mutants of GABA_A_ receptor, *Rdl*, choline acetyltransferase, *Cha*, and electrical synapse, *ShakB*. Instantaneous firing frequency (left) and sample population statistics (right) in indicated genotypes. (Ai) Representative instantaneous firing frequency plots during DD. (Aii) Box plots of peak instantaneous firing frequency. (Aiii) Time to peak frequency in the respective mutants. (Bi) Sample plots of instantaneous firing frequency during flight in the respective mutants. (Bii) Box plots of mean instantaneous firing frequency. (Biii) Minimum frequency for the respective synaptic transmission mutants. (Ci) Sample plot of instantaneous firing frequency during grooming in the respective mutants. (Cii) Mean and (Ciii) Maximum instantaneous firing frequencies during grooming. Boxes represents the 25^th^, 50^th^, and 75^th^ percentiles while whiskers indicate 5^th^ and 95^th^ percentiles of the distribution. Arrowheads indicate distribution mean. Sample sizes as indicated. (**p < 0.01, ***p < 0.001, Mutant vs. WT, Student’s T-test)

Compared to the clear effects on DD spike trains, the flight and grooming pattern generators showed much higher resilience to the same perturbations of the above transmission systems. During flight, only *Rdl* mutants displayed a modest reduction in firing frequency compared to WT counterparts (mean ISI^−1^ of 5.9 Hz vs 8.2 Hz respectively, Figure 5B). For grooming spike patterns, which is intrinsically highly variable, we did not detect significant differences among the mutants tested (Figure 5C), with average mean ISI^−1^ ranging from 0.39 to 1.22 Hz across genotypes, and maximum ISI^−1^ observed between 92 and 120 Hz.

Given the clear alterations in ISI^−1^ profiles of DD spike trains compared to the other motor patterns, we further characterized the impacts of *Rdl, Cha* and *ShakB* mutations on other parameters of ECS discharge repertoire (Figure 6A, cf. [22]). Specifically we examined the stimulation threshold to trigger DD, the onset timing of DD, its duration, and the duration of transmission failure along the GF pathway following electroconvulsive stimulation. While WT and *Rdl* displayed similar thresholds and DD durations (Figure 6B-C), *Cha* and *ShakB* were apparently hypoexcitable as indicated by significantly higher ECS thresholds, and the reduced DD duration (mean durations of 10.0 and 9.4 s respectively) than WT and *Rdl* (13.3 and 11.8 s, Figure 6C). Notably, only *Cha* affected the onset time of DD (22.7 vs 18.3 s for WT, Figure 6D), which has been seen only in a subset of hyperexcitability mutants [22]. In contrast, all three mutants examined displayed prolonged periods of GF failure upon electroconvulsive stimulation, as determined by DLM failure to respond to 1-Hz brain stimuli (Figure 6A, cf. [22]). Modest increases in DLM failure was observed in *Rdl* and *Cha* (36.8 and 36.0 vs 26.0 s for WT) and extreme lengthening in *ShakB* (> 80 s, Figure 6E), a powerful demonstration for the important role of electric synapses in the transmission along the GF pathway that mediate the escape reflex [42, 34, 33, 68].

**Figure 6.**
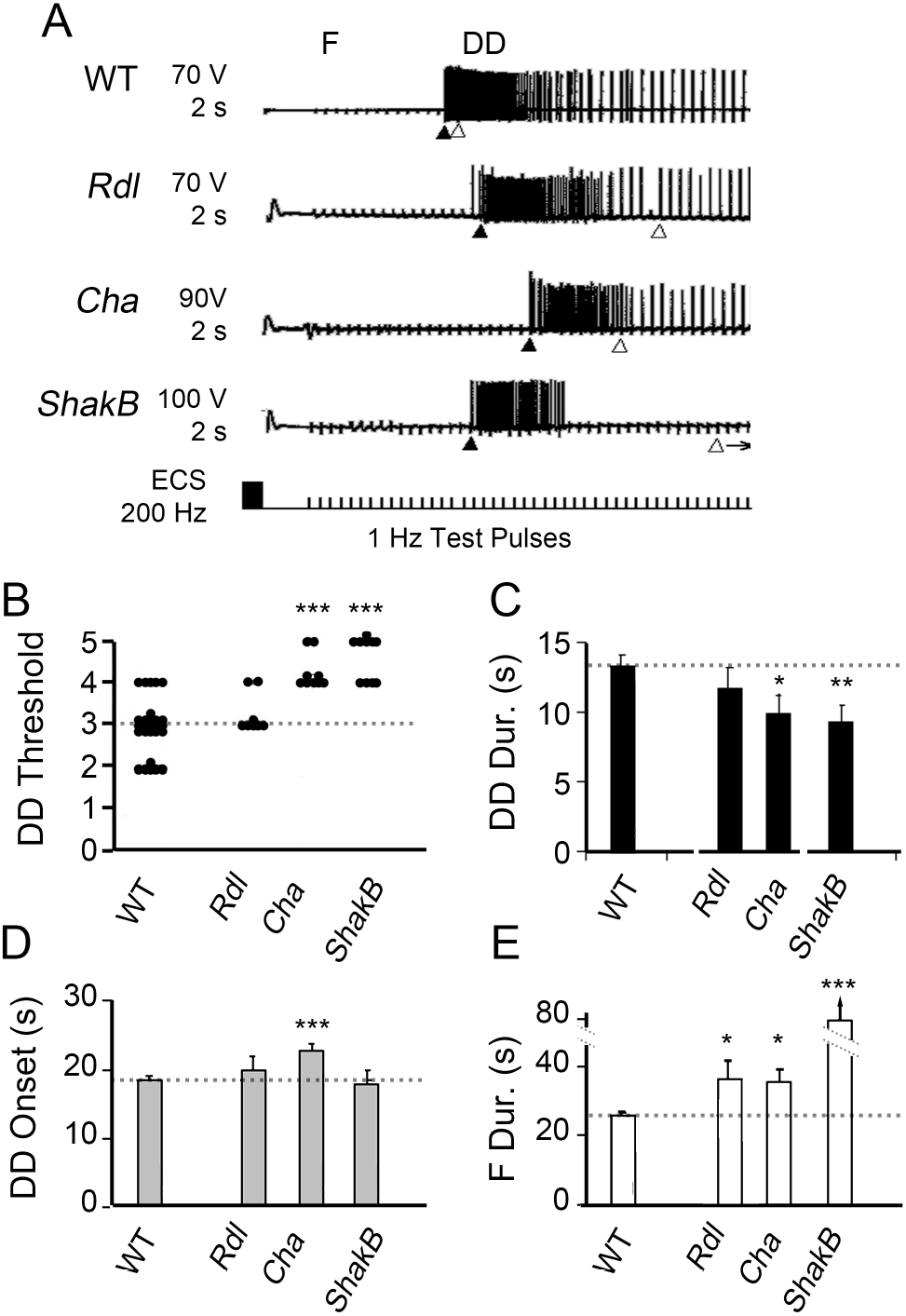
Electroconvulsive stimulation-evoked seizures in WT and *Cha, Rdl*, and *ShakB* mutants. (A) Representative traces of electrical stimulation-evoked GF failure (F) followed by DD in the respective mutants. The onsets of DD (closed triangle) and recovery of GF transmission (open triangle) are indicated below each trace. The stimulation intensity and duration (indicated) in these sample traces were significantly higher than the threshold to trigger DD patterns at saturation levels (Lee & Wu, 2002). (B) Threshold of stimulation required to trigger DD in respective mutants. Increments in values correspond to approximate doubling ECS intensity (1: 50 V, 0.5 s; 2: 50 V, 1 s, 3: 100 V 1 s; 4: 100 V, 2 s; 5: 100 V, 4 s; cf. Lee & Wu 2006). (C) Duration of DD evoked in respective mutants. (D) Elapsed time between ECS and DD onset. (E) Elapsed time between ECS and GF transmission recovery from failure (F Duration) in mutants examined. Note that in *ShakB* GF transmission often failed throughout the entire recording session (lasting > 80 s). For B-E *p < 0.05, ** p < 0.01, ***p < 0.001, Mutant vs WT values, Student’s T-test.

### Poincaré Plots and CV_2_-ISI^−1^ Maps as Distinct Signatures for Electoconvulsive Seizure Spike Patterning in Synaptic Transmission Mutations

The diverse range of alterations to delayed seizure discharges evident in the above analyses (Figures 5 & 6) is corroborated by characteristic differences further revealed through Poincaré plots and ISI^−1^-CV_2_ trajectories of DD spike trains (cf. Figure 4) in neurotransmission system mutants (Figure 7). We found the reduction in peak firing rate of DD in *Rdl* mutants (Figure 6A) was accompanied by an increase in irregular firing, indicated by relatively large deviations from the lower-left to upper-right diagonal of Poincaré plot (Figure 7A). The same spike trains plotted on ISI^−1^ – CV_2_ phase trajectory (cf. Figure 4B) also illustrate more variable trajectories, lacking a general characteristic trajectory compared to WT (Figure 7B). In contrast, *Cha* mutants showed somewhat less irregular firing and ISI^−1^ vs CV_2_ trajectories. However, both *Rdl* and *Cha* DD trajectories appeared to be confined to region with a modest “left-shift” compared to WT, consistent with decreased ISI^−1^ values described above (cf. Figure 5A). Although *ShakB* displayed a similar peak ISI^−1^ value compared to WT flies (Figure 5A), the Poincaré plot indicates more erratic variability in firing frequency within DDs (Figure 7A), which is also illustrated in the ISI^−1^ vs CV_2_ trajectories, with many trajectories scattered over a wide range (Figure 7B). However, the average trajectory suggests similar firing frequency range even though *ShakB* displayed shorter trajectories compared to WT (Figure 7B).

**Figure 7.**
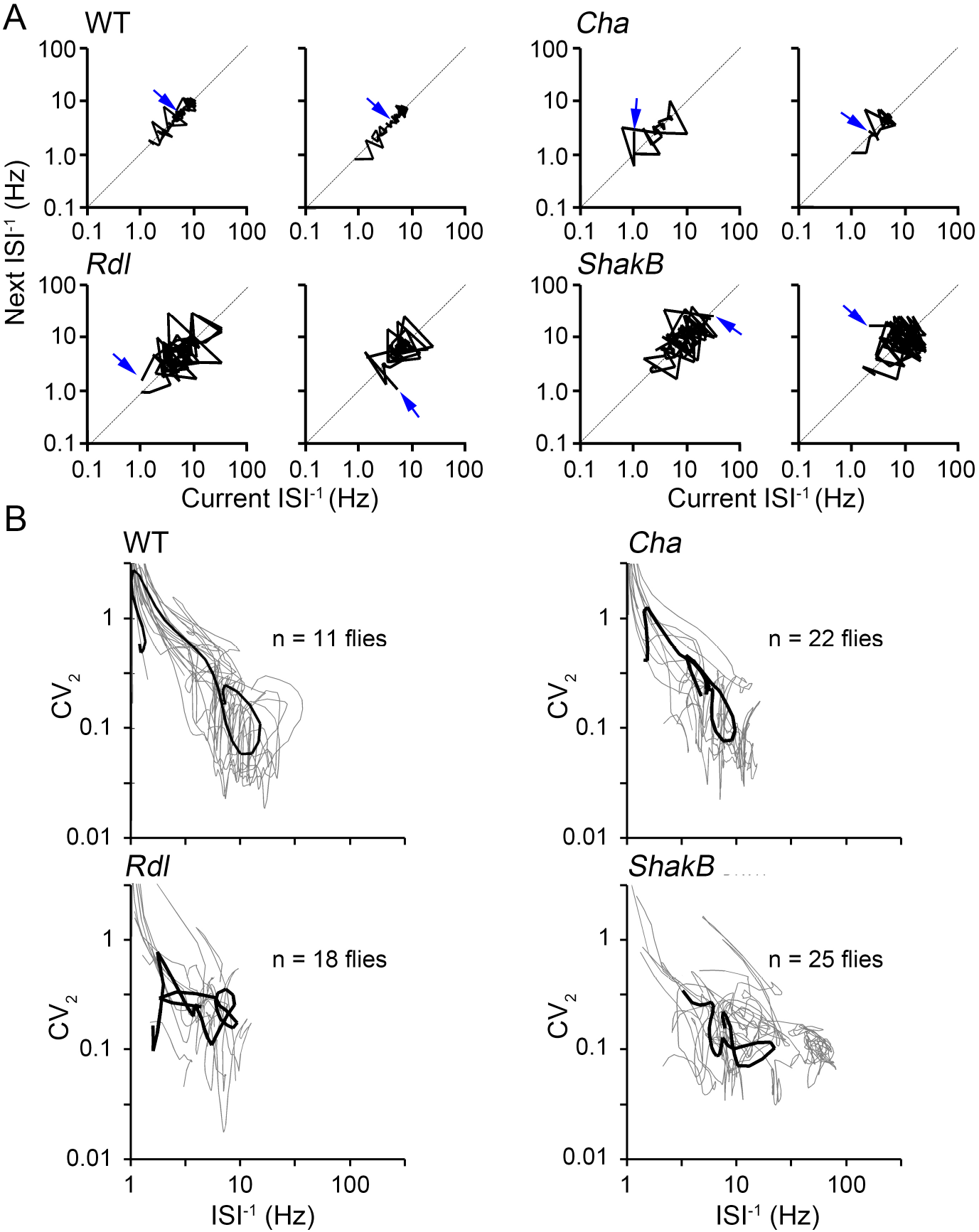
Poincaré plots and ISI^−1^ versus CV_2_ trajectories of DD in synaptic transmission system mutants. (A) Representative Poincaré plots of DD spike trains in WT, *Rdl, Cha* and *ShakB* mutants. Arrow indicates initial interinterval of the spike train. (B) ISI^−1^ versus CV_2_ trajectories from DDs in 10 representative individuals (grey lines) from sample population and averaged trajectory (dark bold line) of DDs across the entire sample population from each geno (sample size as indicated).

### Picrotoxin Blockade of Inhibitory GABA_A_ Receptors: Evolving Seizure Activity Patterns from Flight-Like Firing to Bursting Discharges

The enhanced GABA_A_ receptor function by the *Rdl^MD-RR^* mutation reduced firing rate and altered DLM spike patterning without detectable changes in ECS threshold (Figures 5 & 6). We examined the effects of suppressing the GABAergic system on motor patterns, including seizure-like behaviors. Ingestion of picrotoxin (PTX), a non-competitive antagonist of ionotropic GABA_A_ receptors [69] results in spontaneous spasms that are accompanied by DLM spike discharges [21]. We extended this previous observation to carry out quantitative comparisons between PTX- and ECS-induced discharges, DD. We found that application of a small drop of PTX solution (1 mM, ~ 0.1 *μ*l) to an incision on the compound eye resulted in abnormal DLM spiking, starting within 10 min of application (Figure 8A). Initially, DLM firing appeared rhythmic, with spiking frequencies somewhat higher than those observed during tethered flight (mean ISI^−1^ ~ 10 – 20 Hz, Figure 8B, cf. Figure 1B). Over the course of 30 minutes, short bursts of spikes emerged (Figure 8A) consistent with the previous report on PTX-fed flies [21]. The ISI^−1^ of spikes within these bursts sometimes exceeded 100 Hz (Figure 8B), and the inter-spike intervals were highly variable. These two modes of PTX-induced spiking activity are referred to as ‘flight-like’ and ‘bursting’ activity respectively below. Importantly, the sequence of PTX-induced DLM spiking was gradual and continuous. Furthermore, distinct from ECS-evoked seizure pattern, a period of paralysis followed by a stereotypic discharge pattern was missing in PTX-induced motor activity, suggesting two different modes of seizure activity.

**Figure 8.**
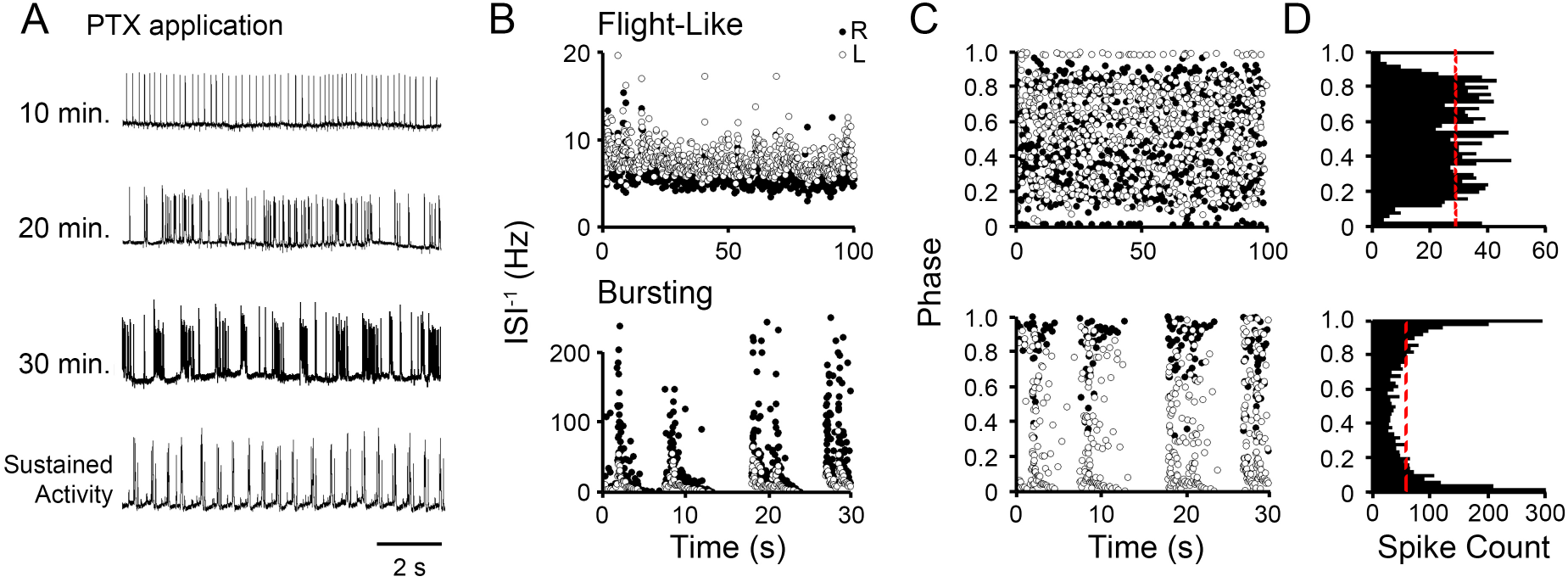
DLM spiking patterns evoked by picrotoxin application. (A) Representative traces of DLM spikes triggered by PTX applied to a punctured eye (see Methods for details) at 10 min intervals over 40 min. Note the transition from rhythmic “flight-like” pattern to repetitive burst discharges. (B) Plots of ISI^−1^ during flight-like and bursting periods from a single individual. Spiking in the left and right DLMs is indicated by open and closed circles respectively. (C) Corresponding phase relations between left and right DLM spiking during the flight-like and bursting states shown in B. (D) Histogram of bilateral phase relations in (C). Note the similarity between the PTX–evoked flight-like and burst firing with respectively the phase relations during flight and grooming shown in Figure 3C-D.

An analysis of the phase relations between bilateral pairs of DLM units revealed striking distinctions between the flight-like and bursting spike patterns induced by PTX (Figure 8C-D). Flight-like firing displayed a relatively uniform phase distribution, with a clear spike-exclusion band approximately ±10% of contralateral spike (Figure 8D), resembling the observations during sustained flight (Figure 3D). Bursting spiking activity, in contrast, displayed a phase-relation histogram with most firing occurring in-phase, resembling the bilateral relationships observed during grooming (Figure 8D, cf. Figure 3D).

In addition to the application of PTX via an incision in the compound eye, we adopted a second procedure, dorsal vessel injection (DV injection, [48]) to investigate whether a global, systemic application of PTX could induce additional spiking patterns. The dorsal vessel serves as a major pulsatile organ which circulates hemolymph [70], and injected solutions circulate throughout the fly within a few seconds. We injected a fixed volume of solution marked with blue #1 dye (0.33 ul) in the tethered fly (Figure 9A, Supplemental Video 2) to ensure the systemic spread of PTX throughout the abdomen thorax and head. We found the two approaches induced the same sequence of spike patterns even though DV injection triggered these events on an accelerated timescale. As Figure 9A shows, PTX injection led to a stereotypic repertoire of a brief wing buzz followed by a ‘wings-up’ posture (see also Supplemental Video 2). Consistent with eye application, PTX injection between 50 and 100 μM induced flight-like DLM spike patterns which evolved into bursting activities within seconds to minutes (Figure 9B). At concentrations of 200 μM or above, we found the flight-like firing pattern was skipped and progression of burst discharge patterns was observed within seconds of injection (Figure 9B).

**Figure 9.**
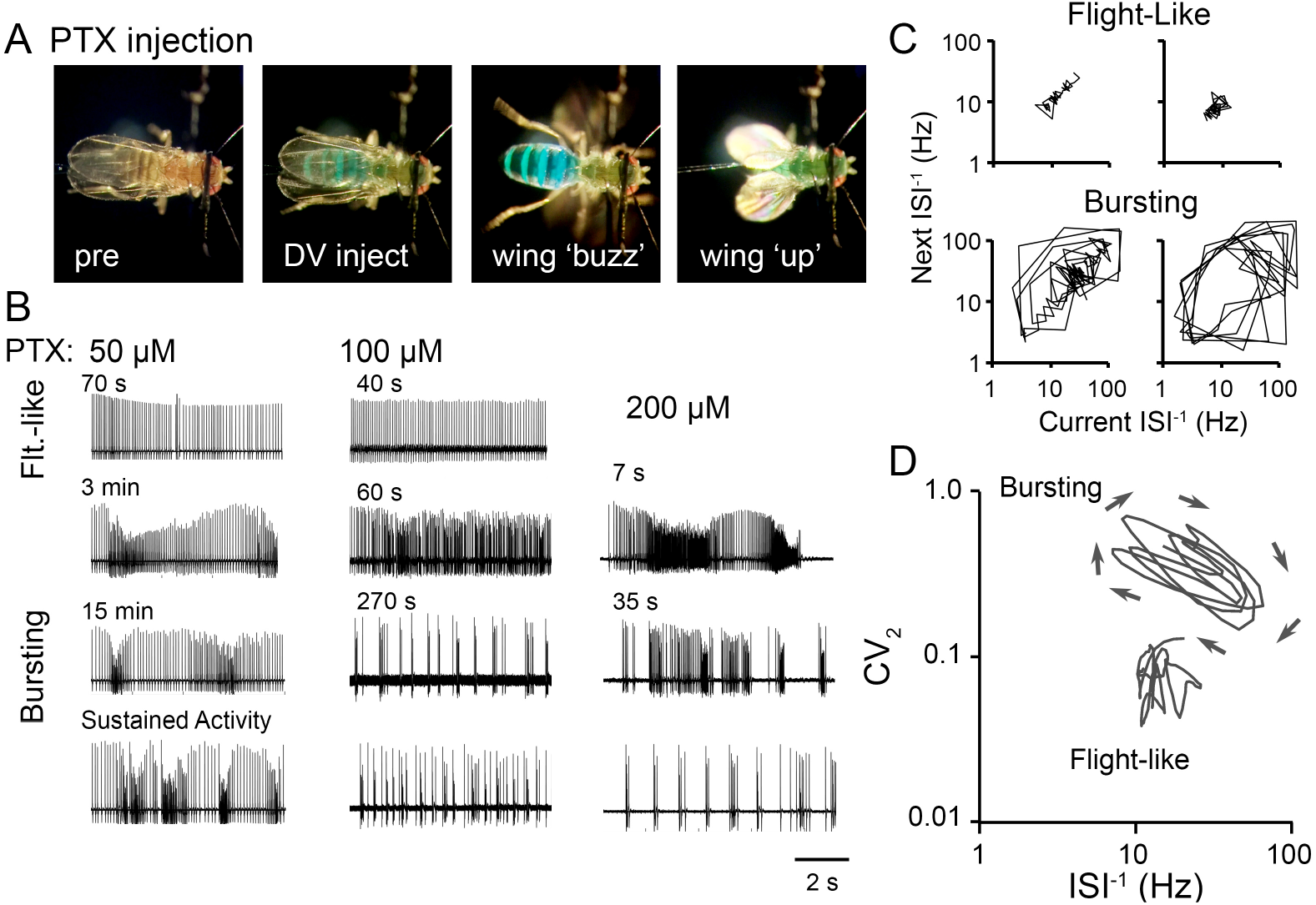
Rapid effects from systemic dorsal vessel (DV) injection of picrotoxin. (A) Video frames illustrating behavior and posture from a fly before and after 100 *μ*M PTX DV injection. A transient wing ‘buzz’ was often observed, and flies displayed a ‘wings-up’ posture for several hours before death. (B) Representative spiking traces from three individual flies injected with either 50, 100 or 200 μM PTX, respectively. Traces illustrate the sequential events, evolving from flight-like to bursting patterns, at different times after injection as indicated. At 50 *μ*M, discrete inter-burst gaps did not fully develop, and at 200 μM, the initial flight-like state was not captured and spike activity (7 s) rapidly evolved into bursting activity. (C) Poincaré plots of DLM spiking displaying flight-like or bursting spike patterns (~5-s) from two representative individuals (100 *μ*M PTX inject). (D) ISI^−1^ vs CV_2_ trajectories from ~ 5-s portions of the spike trains shown in C during the flight-like and bursting DLM spiking phases.

The two categories of PTX-induced spike patterns are clearly contrasted by their trajectories in Poincaré plots (Figure 9C). The flight-like spike patterns displayed PTs which were relatively compact, with occasionally sharp deviations, corresponding to spike doublets or brief gaps in firing, which were notably absent during sustained flight in wild-type flies (compare to Figure 4A). Bursting spike patterns displayed strikingly different PTs, characterized by stereotypic recurrent sequences of successive ISI^−1^ values, corresponding to recursive bursts. Plots of the ISI^−1^ vs. CV_2_ served to further contrast these features (Figure 9D). PTX-induced flight-like firing displaying a more stable phase trajectory with CV2 ~0.1, and ISI^−1^ higher than flight (~10 – 20 Hz), whereas bursting appeared as a looping trajectory. Each burst cycle started with at a relatively lower ISI^−1^ and high CV_2_, then accelerated to high ISI^−1^ and lower CV_2_, corresponding to the long inter-burst interval followed by the fast peaking and slow decay of the firing frequency within the burst.

## Discussion

Due to the ease of electrical monitoring, coupled with the tractable genetic manipulations in *Drosophila*, the indirect flight muscles, DLMs, have been frequently used for studying consequences of mutations of identified genes on membrane excitability and synaptic transmission (for early examples, see [71, 19, 72, 73]. Mutational analyses of a number of motor programs that drive DLM activity have also been carried out. DLM responses have enabled characterization of the GF pathway-mediated jump-and-flight escape reflex triggered by visual or vibrational stimulation [41, 34, 35, 74, 68, 75, 76, 77] which leads to escape-related wing depression [33, 78]. Additionally, DLM recording has enabled detailed genetic analyses of the pattern generators that drive flight [79, 80, 27], grooming [34, 81, 57], and electroconvulsively induced seizure discharges, including the initial discharge, ID, and delayed discharge, DD have also been carried out [20, 82, 23, 21, 22, 83].

In the present study, direct comparisons of spike patterning aim at an initial qualitative and quantitative delineations among three distinct modes of DLM spiking associated with DD, flight, and grooming motor programs.

### DLM-Activity Signatures Associated with Seizures, Flight and Grooming Motor Patterns

The spiking activity of the six DLMs on each side of the fly thorax is driven by five motor neurons in the mesothoracic ganglion, with the lower four muscles (DLMc-f) each innervated by a single motor neuron (MN1 – 4) on the ipsilateral side, while the top two muscles (DLMa and DLMb) are innervated by a single contralateral motor neuron, MN5 [58, 84, 85]. The MN5 is particularly well characterized, in terms of neurite outgrowth [86, 87, 88] as well as excitability [89]. Individual Ca^2+^ [90] and K^+^ [91] currents within the MN5 soma have been characterized, and their contributions to action potential generation computationally modeled [92]. Notably, the MN5 and the DLMa-b fibers are capable of following direct stimulation one-to-one well beyond 100 Hz [73, 34]. Thus, recording from DLMa or DLMb faithfully registers the MN5 activity, facilitating the analyses of a variety of motor patterns spanning a wide range of firing frequencies.

To quantitatively differentiate DLM spiking patterns associated seizure discharges, flight and grooming activities (Figure 1), we characterized several parameters to capture salient features of spike firing frequency (Figure 2), spike timing relationships between contralateral units (Figure 3), and the evolving trajectory of varying firing regularity and frequency during episodes of spiking activity (Figure 4). The instantaneous firing frequency, ISI^−1^, has been extensively utilized to quantify DLM spiking frequency (e.g. [93, 53]), and our analysis revealed distinctive ISI^−1^ distributions during the three respective motor programs. Further, the spike timing relationships between the left and right units (Figure 3), as well as temporal characteristics revealed by Poincaré plots and ISI^−1^-CV_2_ trajectories clearly separate these firing patterns into distinct categories (Figure 4).

Consistent with previous reports [29, 53], we noted that the left-right phase distribution of flight spike trains displayed “exclusion band” in which synchronous firing between the bilateral DLM pair is very rare. In contrast, during DD spiking no discernable phase relation in timing between the left and right units was observed, while grooming spikes tended to be phase-correlated (Figure 3). These findings suggest that the exclusion band is specific to flight motor patterns, although the precise cellular and molecular mechanisms remain to be uncovered. Indeed, electrical coupling between motor neurons [53], or inhibitory glutamate channels (gluCl[94]) have been proposed to mediate this mutual feedback inhibition.

Among DD, flight and grooming, a readily apparent qualitative distinction is the ISI variation in their spike patterns, with low variability associated with flight, high variability with grooming, and DD in between (Figure 1). To contrast the spike interval variability associated with the three motor patterns, we employed two related approaches to analyze spike trains, Poincaré plots (Figure 4A) and of the ISI^−1^ vs CV_2_ plots (Figure 4B). Poincaré plots have been frequently utilized in characterizing variability in heart beats (e.g. [95, 96]) and in neuronal action potentials (e.g. [97]). We found that Poincaré plots succinctly display the differences in spike interval variability among the three types of spike patterns. Because the Poincaré trajectories from individual spike trains cannot be readily combined to produce an ‘average’ ensemble trajectory, we utilized the instantaneous coefficient of variation CV_2_ ([52], Figure 4B) to compile the merged, ensemble spike interval variability across a collection of spike trains that presented an overall signature of the temporal progression of spike interval variability for each motor program. It should be noted that CV_2_ documents a concise measure of variability of inter-spike intervals (e.g. [98, 99]), and can be derived from the Poincaré trajectory. Indeed, plots of the ISI^−1^ vs CV2 trajectory of spike trains captured the salient features of the sustained rhythmic firing (low CV2) in flight spike trains, brief bouts of high-frequency firing during grooming (high CV_2_, variable ISI^−1^), and the characteristic progression of firing frequency and spike interval variability during delayed seizure discharges (Figure 10).

**Figure 10.**
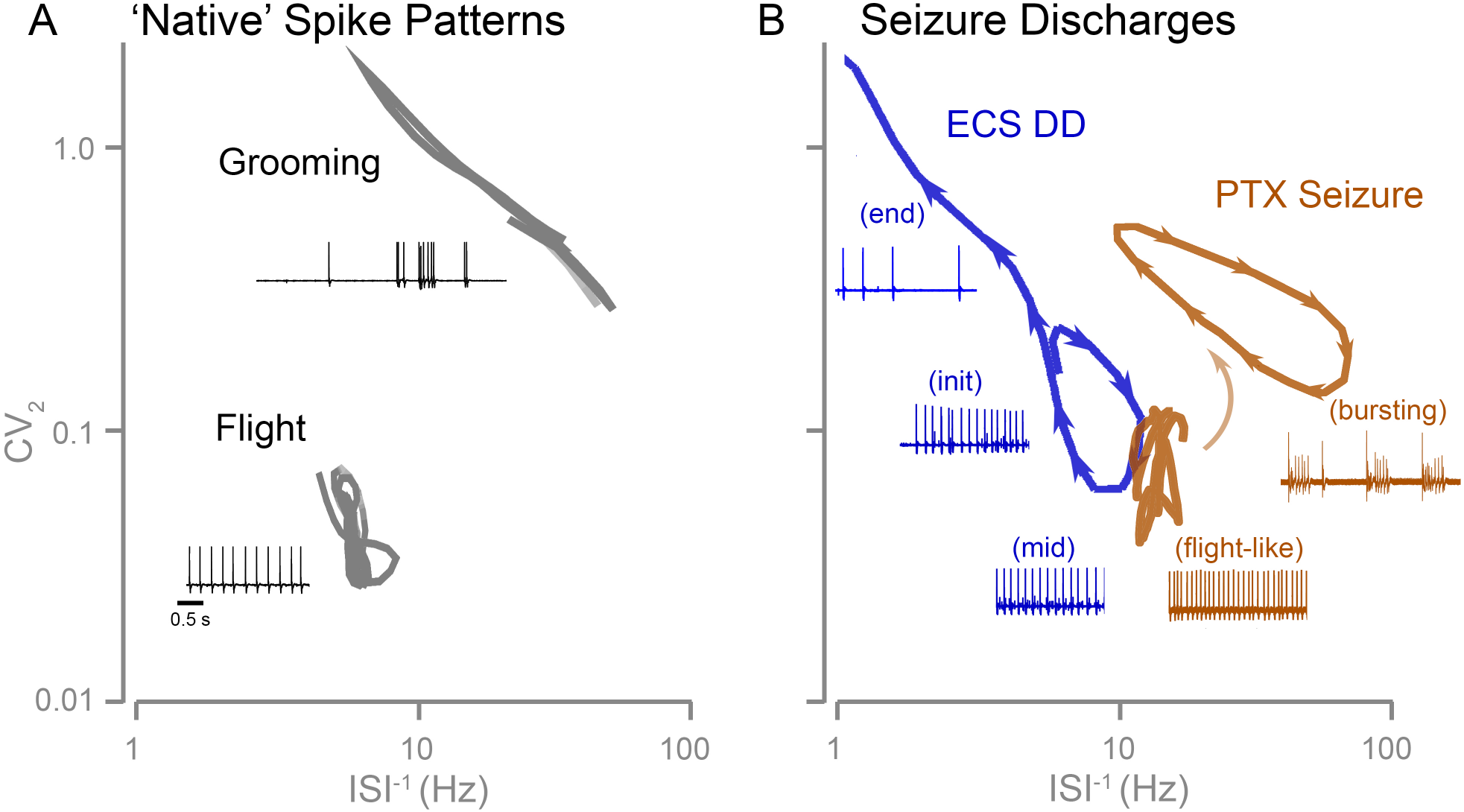
ISI^−1^ vs CV_2_ plots reveal distinctions in DLM spike patterns during flight, grooming and seizure discharges. Summary of averaged ISI^−1^ vs CV_2_ trajectories of DLM spiking during different motor patterns. Insets: Representative traces illustrating different spiking activities associated with the distinct trajectories occupying different regions of the ISI^−1^-CV_2_ plot. (A) Trajectories of ‘native’ spike patterns during flight and grooming. (B) Trajectories of electroconvulsive stimulation- and PTX-evoked seizure spike patterns. Arrow heads indicated trajectory direction during seizure discharges.

### Neurotransmitter Systems in Shaping Seizure Discharges and Native Motor Patterns

Despite their obvious importance, a more comprehensive characterization remains to be accomplished to reveal the involvement of individual neurotransmitter systems in generating flight, grooming, and ECS-related activity in *Drosophila*. We aimed to provide an initial glimpse of the general vulnerability of the respective motor patterns to perturbations of excitatory, inhibitory and electrical transmission systems. The results also provide a context for the modulatory role of biogenic amines on these motor programs that have been examined to some depth [100, 80, 101]. We selected mutant alleles with well-described perturbations of neurotransmission. Specifically the *Cha* and *ShakB* flies displayed clear disruptions of cholinergic and electrical transmission respectively along GF pathway [41, 42], a robust neuronal circuit critical for escape behaviors [32, 33, 35, 36, 78]. For the *Rdl* mutant, the insecticide resistance phenotype has led to identification of the GABA_A_ receptor gene and this particular allele that has been extensively studied [67]. Studies in cultured neurons have demonstrated altered sensitivity to GABA_A_ blockade [65, 102].

Given the importance of these synaptic transmission systems on activity in the nervous system, it was somewhat unexpected that relatively minor modifications to flight and grooming patterning were uncovered (Figure 5B-C). Indeed, we observed that these mutant flies were capable of flight and grooming. Due to the intrinsically high variability across grooming spike trains, it is difficult to conclusively quantify any minor differences caused by the mutations. During flight, however, it was clear that only *Rdl* mutants displayed a reduction in DLM firing frequency, supporting the hypothesis of direct feedback regulation among DLM motor neurons underlying flight patterning [29, 53]. Conceivably, such feedback interactions may be mediated via GABAergic interneurons.

ECS discharges, in contrast to flight and grooming motor activities, appeared to be quite sensitive to disruption of neurotransmission systems. Interestingly, perturbations to GABAergic, cholinergic and electric synaptic systems each led to alterations in overlapping but not identical subsets of DD parameters (Figures 5-7). We found that enhancing GABAergic effectiveness in the *Rdl* allele and suppressing cholinergic transmission in *Cha* mutations severely reduced the peak firing frequency of delayed seizure discharges (Figure 5A), while *Cha* and the gap junction mutant *ShakB* displayed increases in the threshold to induce a DD seizure with a corresponding shortened DD duration. Consistent with their integral role of transmission along the GF pathway, mutations in these transmission systems also prolonged the period of circuit failure following the high-frequency electroconvulsion stimulation. In particular, none of the *ShakB* mutants recovered over the > 80 s recording period (Figure 6). Notably, our analysis of DD in *ShakB* corroborates findings of a previous report of increased seizure thresholds in the same mutant alleles (Song & Tanouye, 2006). The alterations in spike patterning during DD were also evident in that *Rdl* and *ShakB* displayed more variable Poincaré plots and distinct ISI^−1^ vs CV_2_ trajectories compared to WT and *Cha* counterparts (Figure 7).

### Distinct Modes of Seizure Discharges Induced by Electroconvulsive Stimulation and GABAergic Blockade

Electroconvulsive stimulation and proconvulsant administration are two means for seizure induction commonly used in both vertebrates and invertebrates [103]. Several studies have utilized the proconvulsant picrotoxin (PTX), a noncompetitive GABAA antagonist, to induce seizure activity in *Drosophila* [21, 104, 105], However, the resulting seizure discharge pattern was reported to be qualitatively distinct from the ECS discharge repertoire [21]. We found that PTX application evoked a stereotypic, evolving DLM activity patterns (Figures 5–6). Unlike DD which occurred over a discrete period (~ 15 s), PTX-evoked activity continued throughout the recording period (> 1 hr), invoking the impression of the release or run-away of certain native, patterned activity. Initially, the DLM displayed a spontaneous, rhythmic spiking pattern reminiscent of flight activity in terms of firing frequency and the phase relation between left and right DLM spiking (compare Figure 4 ‘flight’ and Figure 8 ‘flight-like’). This mode of activity was succeeded by bursts of spikes with instantaneous firing frequencies exceeding 100 Hz during which spiking was generally synchronized between the left and right sides. Loss of central inhibition is known to release a number of motor programs in insects [106], with a notable example of decapitation-triggered courtship motor programs in preying mantids [107]. Therefore, it is tempting to speculate that automatisms observed in the present study during ECS- and PTX-induced seizures may represent two separate forms of synchronized activities ‘released’ by specific treatments [108, 109, 110].

As summarized in Figure 10B, the ISI^−1^ vs. CV_2_ plot of the spike trains elucidates distinctions between DD and PTX-evoked activity indicating separate neurophysiological dysfunctions. The distinction between electrically and pharmacologically triggered discharges has also been noted in vertebrate seizure models. Indeed, the expression of seizure-like behavioral sequence evoked by pentylenetetrazole (a GABA_A_ receptor antagonist) is distinct from seizure discharges evoked by maximal electroconvulsive stimulation in rats [111, 112].

These results serve to illustrate the importance to provide precise descriptions of seizure induction methods to avoid confusion in the literature when studying the molecular and cellular mechanisms in generating motor phenotypes. Furthermore, in future neurogenetic studies of ‘seizure’ behavior in *Drosophila*, a combination of pharmacological and electrical seizure induction methods can enhance the latitude and depth of analysis for the roles of individual genes participating in distinct mechanisms underpinning seizure activity.

Our study also demonstrates that in conjunction with spike firing rate and frequency distribution analyses, Poincaré plots adequately characterize the ISI variation along individual spike trains. The stochastic nature of spike trains, with ISI sequences varying from trial to trial within the same fly and between individuals, some inherent features characteristic of a particular motor pattern may be masked or overlooked. Nevertheless, the ability to construct CV2 for individual spike trains, which allows to derive an averaged ensemble trajectory pooled from a population flies, may serve as signatures in the ISI^−1^ vs. CV_2_ plot for the various motor patterns and to delineate how they are modified by genetic or pharmacological manipulations.

## Acknowledgments

We thank members of the Wu Lab for their helpful comments over the course of this project. This work was supported by NIH grants: GM080255, GM088804, AG047612, and NS098590 to CFW, and NS082001 to AI

