## Supplementary figures and images for "Distinctions among electroconvulsion- and proconvulsant-induced seizure discharges and native motor patterns during flight and grooming: Quantitative spike pattern analysis in *Drosophila* flight muscles"

### Supplemental Figure 1

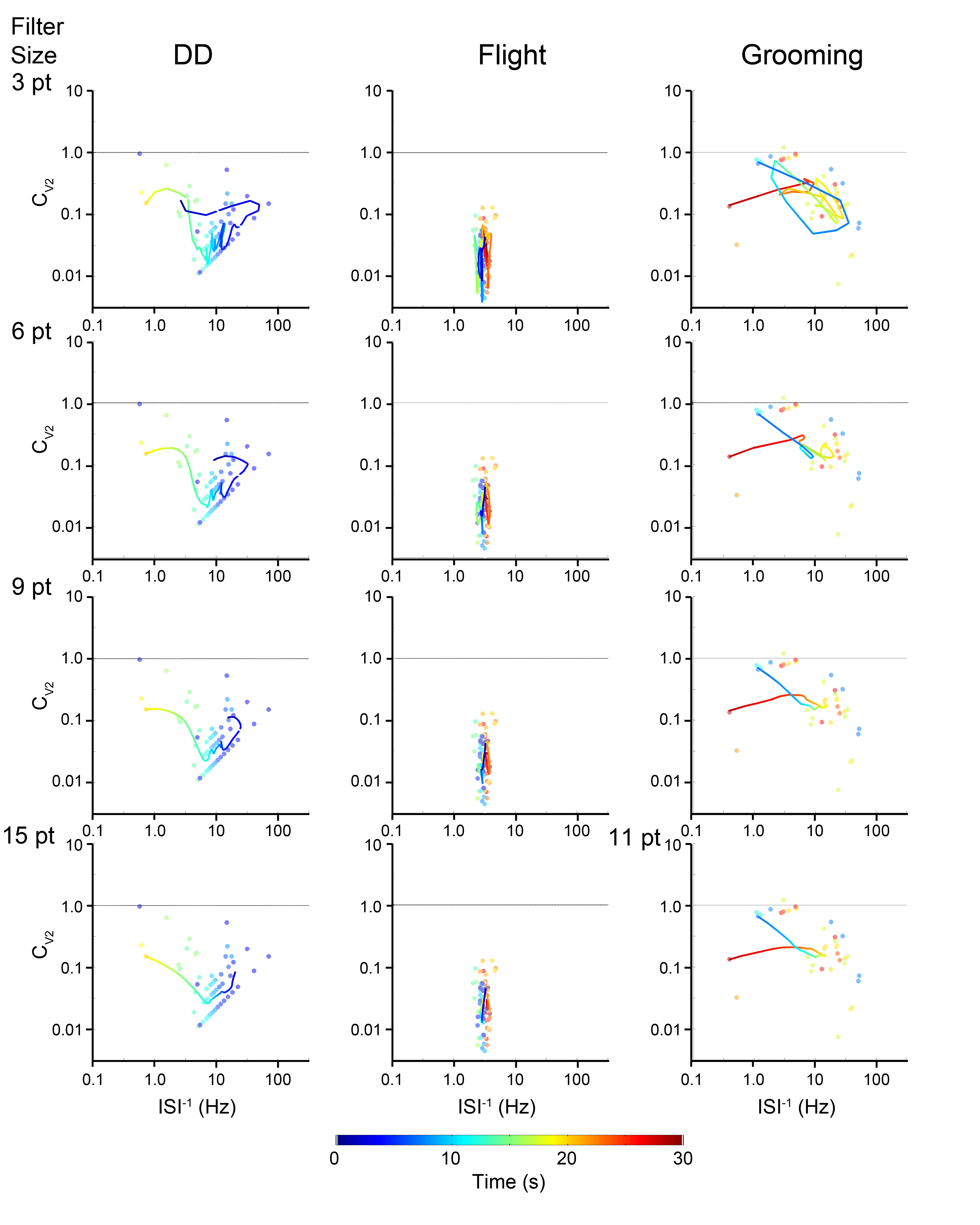
